# rphylopic: An R package for fetching, transforming, and visualising PhyloPic silhouettes

**DOI:** 10.1101/2023.06.22.546191

**Authors:** William Gearty, Lewis A. Jones

## Abstract

1. Data visualisation is vital for data exploration, analysis, and communication in research. Moreover, it can bridge gaps between researchers and the general public by making research findings more accessible and engaging. Today, researchers increasingly conduct their data analyses in programming languages such as R and Python. The availability of data visualisation tools within these environments supports the generation of reproducible data analyses and visualisation workflows. However, resources from external databases are frequently required for data visualisation necessitating integration between these databases and the platforms in which researchers conduct their analyses.
2. Here, we introduce rphylopic, an R package for fetching, transforming, and visualising silhouettes of organisms from the PhyloPic database. In addition to making over 7,000 organism silhouettes available within the R programming language, rphylopic empowers users to modify the appearance of these silhouettes for ultimate customisability when coding production-quality visualisations in both base R and ggplot2 workflows.
3. In this work, we provide details about how the package can be installed, its implementation, and potential use cases. For the latter, we showcase three examples across the ecology and evolutionary biology spectrum.
4. Our hope is that rphylopic will make it easier for biologists to develop more accessible and engaging data visualisations by making external resources readily accessible, customisable, and usable within R. In turn, by integrating into existing workflows, rphylopic helps to ensure that science is reproducible and accessible.

## Introduction

The world is now in an age of big data, and the biological sciences are no exception (Spengler, 2000). With the advent and growth of online databases such as the Global Biodiversity Information Facility, Ocean Biogeographic Information System, the Paleobiology Database, and GenBank, biologists are now flooded with ecological, evolutionary, and conservation data. Alongside the growth of these databases, advanced computational methods to analyse these data have been developed. However, despite–and oftentimes because of–these advancements in analytical techniques, data visualisation remains a key component of any scientific data analysis workflow (Ali et al., 2016). Specifically, data visualisation is vital for the summarisation and dissemination of complex ideas and results (Ware, 2019). Moreover, data visualisation can play a critical role in communicating research findings to stakeholders (Pauwels, 2006). However, as theories and findings become more complex, effectively visualising data becomes increasingly difficult (Ali et al., 2016; Midway, 2020).

Fortunately, numerous tools have recently been developed to perform effective and efficient data visualisation (Caldarola & Rinaldi, 2017). While many of these tools are platform-or language-specific, such as the popular R package ggplot2 (Wickham, 2016), others are platform-agnostic. PhyloPic (https://www.phylopic.org/) is an online open-access database of silhouette images of animals, plants, and other life forms. To date, the silhouettes have been created by over 500 volunteer contributors, are available for reuse under various Creative Commons licenses, and are stored within a taxonomic/phylogenetic framework (e.g. phylum Mollusca and clade pan-Mollusca are both valid). While PhyloPic is almost never cited within scientific publications, silhouettes from the database have been utilised to effectively and simply convey taxonomic information for innumerable scientific articles, conference contributions, and much more. However, despite this already wide array of applications, usage hurdles exist due to its platform-agnostic nature. For example, while silhouettes can be downloaded in raster or vector formats, these can still be difficult to integrate into a coding-based scientific workflow, especially when the user is trying to maintain reproducibility. Often, custom code and/or post-production editing are required to accommodate these images.

Here we describe rphylopic, a dedicated R package for interfacing with the PhyloPic database. The package contains functions for accessing, transforming, and visualising PhyloPic silhouettes. We first provide instructions for installing the package and details on its implementation. Second, we demonstrate the functionality available in rphylopic through usage examples. Finally, we provide details about the resources we have made available to the community and to support rphylopic users.

## Installation

The rphylopic package can be installed from CRAN using the `install.packages` function in R (R Core Team, 2023):

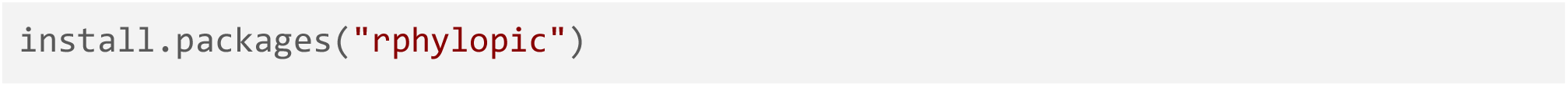

If preferred, the development version of rphylopic can be installed from GitHub via the remotes R package (Csárdi et al., 2021):

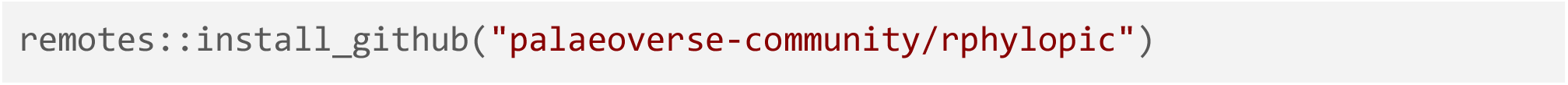

Following installation, rphylopic can be loaded via the `library` function in R:

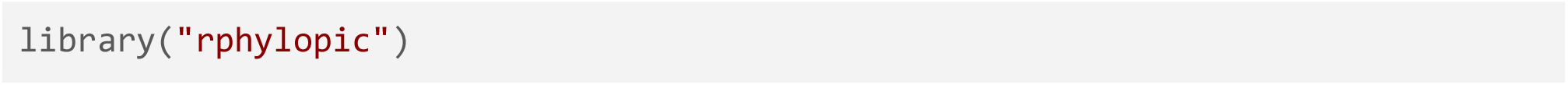

### Dependencies

The rphylopic package depends on R (> 4.0) (R Core Team, 2023) and imports functions from: curl (Ooms, 2023), ggplot2 (Wickham, 2016), graphics (R Core Team, 2023), grid (R Core Team, 2023), grImport2 (Potter & Murrell, 2019), httr (Wickham, 2023), jsonlite (Ooms, 2014), methods (R Core Team, 2023), pbapply (Solymos & Zawadzki, 2023), png (Urbanek, 2022), and rsvg (Ooms, 2022). The package was developed with the support of the R packages devtools (Wickham, Hester, et al., 2022), testthat (Wickham, 2011), and roxygen2 (Wickham, Danenberg, et al., 2022).

## Implementation

A summary of the functions currently available in rphylopic is provided in Table 1. Detailed description is provided below followed by usage examples to demonstrate the functionality and versatility of the package.

**Table 1.** A summary table of the functions currently available in the rphylopic R package.

| Function | Description |
| --- | --- |
| add_phylopic | Adds PhyloPic silhouette(s) layer to ggplot2 plots |
| add_phylopic_base | Adds PhyloPic silhouette(s) to base R plots |
| flip_phylopic | Flips a PhyloPic silhouette along a horizontal or vertical axis |
| geom_phylopic | Plots data points as PhyloPic silhouette(s) for ggplot2 |
| get_attribution | Gets the attribution data for a given PhyloPic silhouette |
| get_phylopic | Gets the image data for a given PhyloPic silhouette |
| get_uuid | Gets the unique identifier for a given PhyloPic silhouette |
| pick_phylopic | Enables visual selection of PhyloPic silhouettes |
| recolor_phylopic | Recolours a given PhyloPic silhouette |
| rotate_phylopic | Rotates a given PhyloPic silhouette |
| save_phylopic | Saves a given PhyloPic silhouette |

### Fetching

Image data (i.e. silhouettes) are fetched from the PhyloPic application programming interface (API) service via the `get_phylopic` function. The function requires an image universally unique identifier (UUID), which can be provided by the user for a known silhouette. If the user does not know the UUID for the desired image, the `get_uuid` function can be used to retrieve the image UUID associated with a specific taxonomic name:

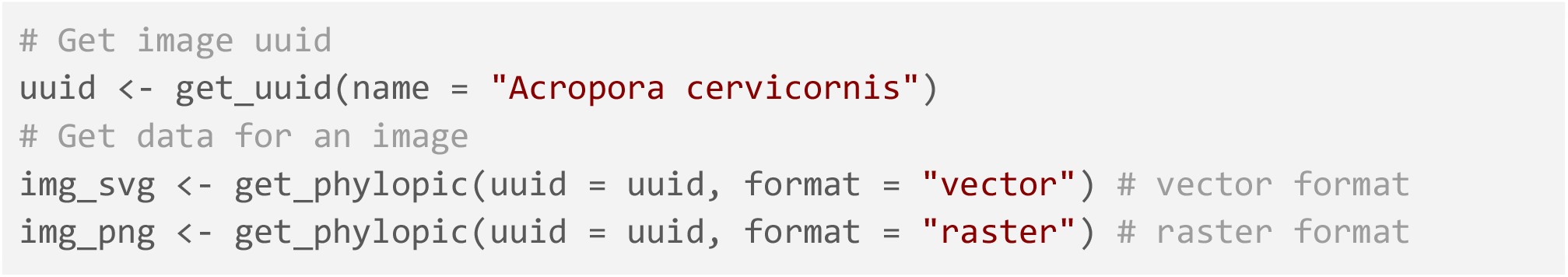

Silhouettes can be returned as both Scalable Vector Graphics (SVG) and Portable Network Graphics (PNG), depending on the user-defined format. An additional argument (`height`) can be defined for the “raster” format to control the image height in pixels.

Multiple silhouettes can exist for a single taxonomic name in PhyloPic, particularly at higher taxonomic levels. To help facilitate user selection from various image silhouettes, we provide the `pick_phylopic` function to enable users to interactively pick an image through visualising available options (Fig. 1):

**Figure 1.**
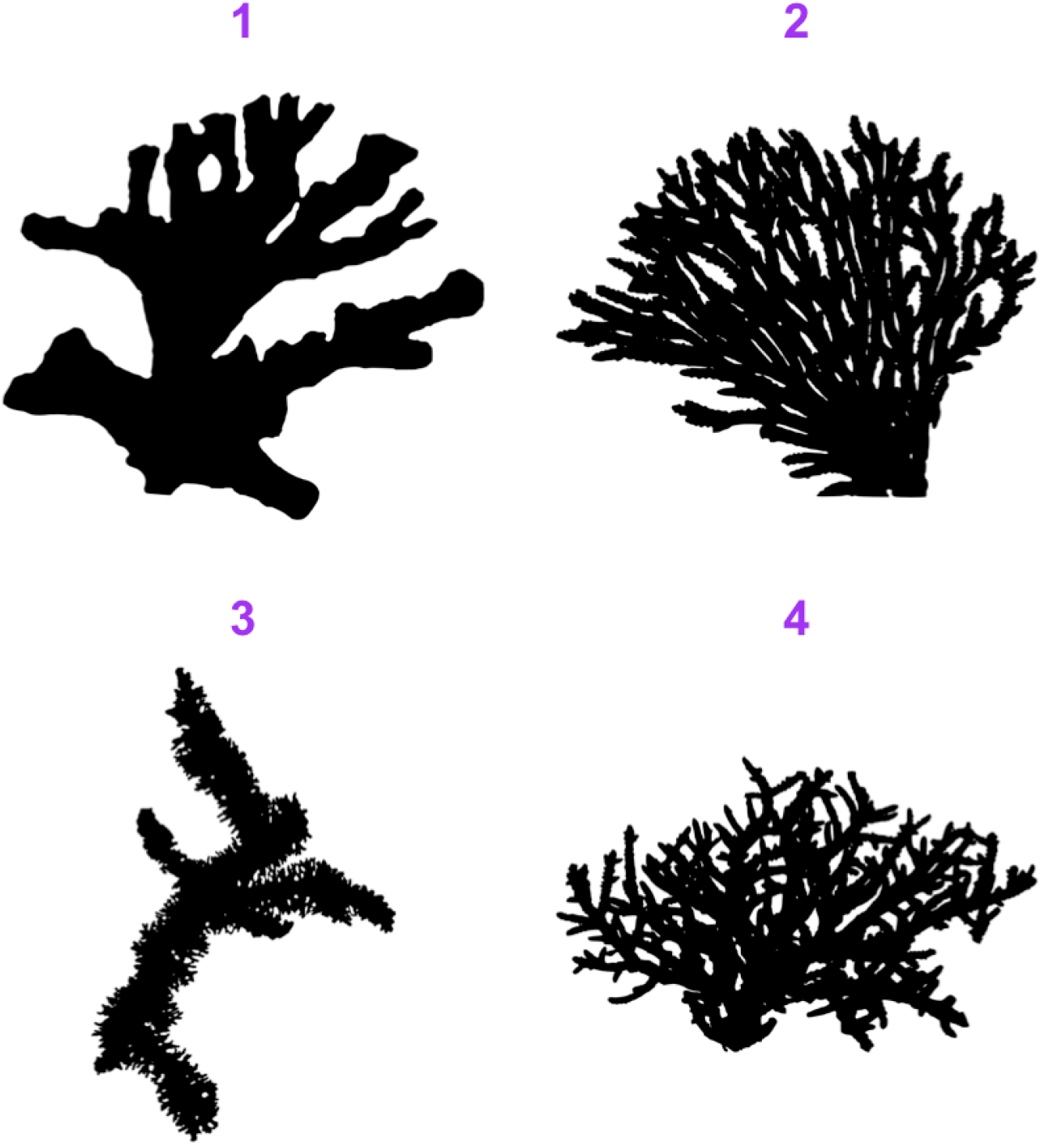
An example of the output from `pick_phylopic` for selecting image silhouettes. The image silhouettes are from PhyloPic (https://www.phylopic.org/; T. Michael Keesey, 2023). Silhouette 1 was contributed by Victor Piñon-González (2023; CC0 1.0) and silhouettes 2–4 were contributed by Guillaume Dera (2022; CC0 1.0).

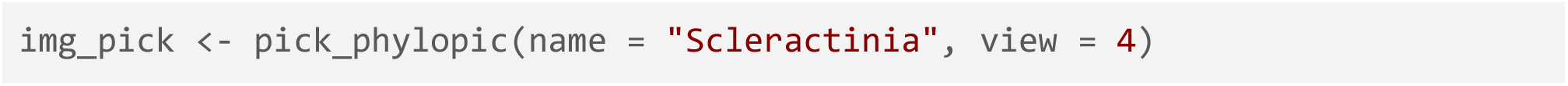

The function `save_phylopic` is provided to allow users to conveniently save any silhouettes in a range of formats, including: Portable Document Format (PDF), Portable Network Graphics (PNG), Scale Vector Graphics (SVG), Tag Image File Format (TIFF), Joint Photographic Experts Group (JPEG), and bitmap (BMP). Note, this is not required if all further transformations and visualisations will be conducted within R; however, we include this function for users who may still need to use these silhouettes outside of R.

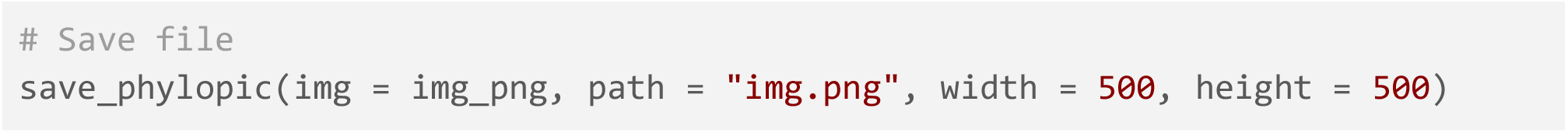

### Transformation

Once a silhouette is picked and saved in the user’s R environment, it may be useful to transform the image to better suit the particular visualisation of interest. We have implemented three user-friendly functions to accommodate three transformations that might be desired by the user: flipping, rotating, and recolouring. All three functions work on both SVG and PNG versions of silhouettes.

The `flip_phylopic` function can be used to flip a silhouette horizontally and/or vertically. This may be useful if, for example, the user wants all of the silhouettes to face the same direction.

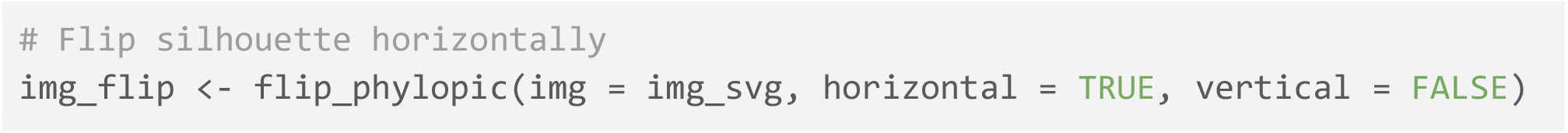

The `rotate_phylopic` function can be used to rotate a silhouette an arbitrary number of degrees. This may be useful when trying to align a silhouette with text or other objects within a figure.

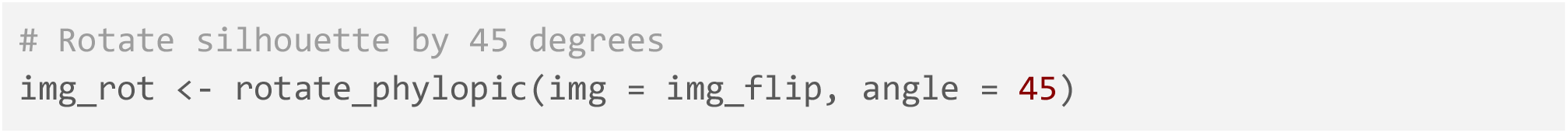

Finally, the `recolor_phylopic` function can be used to modify the colour and/or transparency of a silhouette. The vast majority of PhyloPic silhouettes are black and fully opaque by default. However, it may be useful to change this when the user is trying to either match an existing visualisation colour palette or trying to convey extra information, such as categorical data, through colour.

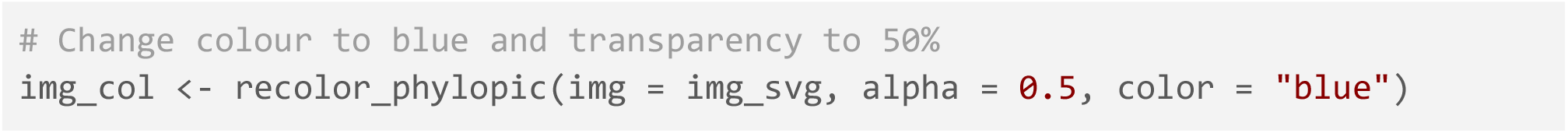

For convenience, we have also included these transformation options within all of the visualisation functions discussed in the next section. However, when the same transformed silhouette will be used for multiple visualisations, we suggest transforming the silhouette first, saving it as a new object, then using this new object for visualisation purposes.

### Visualisation

In recognition of the wide range of users that may use the package, we have included functions to add the desired PhyloPic silhouettes to both base R and ggplot2 plots. Further, for convenience, each of the following functions works with the names of species/clades (the `name` argument/aesthetic), UUIDs (the `uuid` argument/aesthetic), or picture objects (SVG or PNG) (the `img` argument/aesthetic). For simplicity, we will only showcase the use of names here.

The `add_phylopic_base` function can be used to add one or more silhouettes to a base R plot. The function behaves very similarly to the base R `points` function, where x and y coordinates are supplied for the centres of the silhouettes. The `ysize` argument can be used to vary the size of the silhouettes. For convenience, if any of these arguments are vectors, the other arguments will be recycled as necessary. For example, a single silhouette can be plotted at multiple x and y coordinates, or multiple silhouettes can be plotted at multiple y coordinates but all with the same x coordinate.

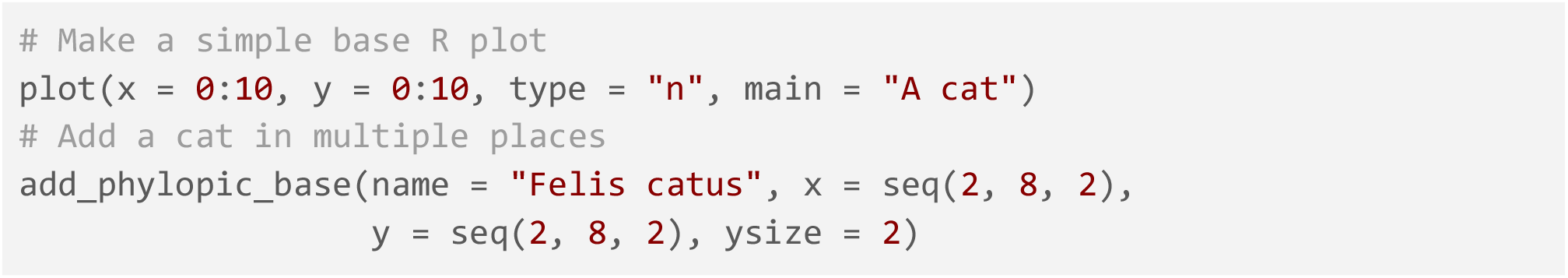

The equivalent function for ggplot2 usage is the `add_phylopic` function, which adds PhyloPic silhouettes as a separate layer to an existing ggplot object. It works very similarly to the `annotate` function in the ggplot2 package (Wickham, 2016). A common use case for this is to add a PhyloPic silhouette as a background image:

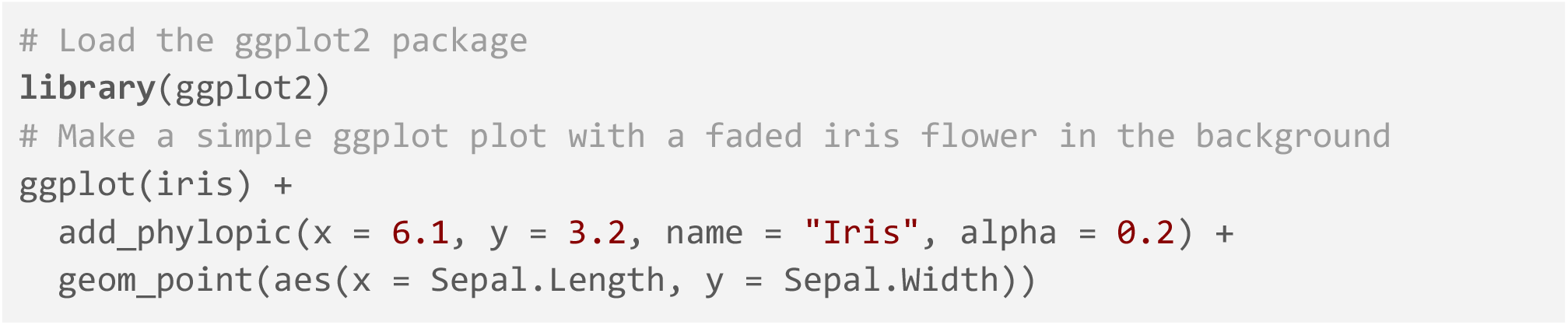

In case the user already has a data frame of data, we have also provided the `geom_phylopic` function, which acts like the `geom_point` function from ggplot2, except the specified silhouettes are used as points. This can be combined with other functions (e.g. `facet_wrap`) from ggplot2 and other packages to create complex visualisations:

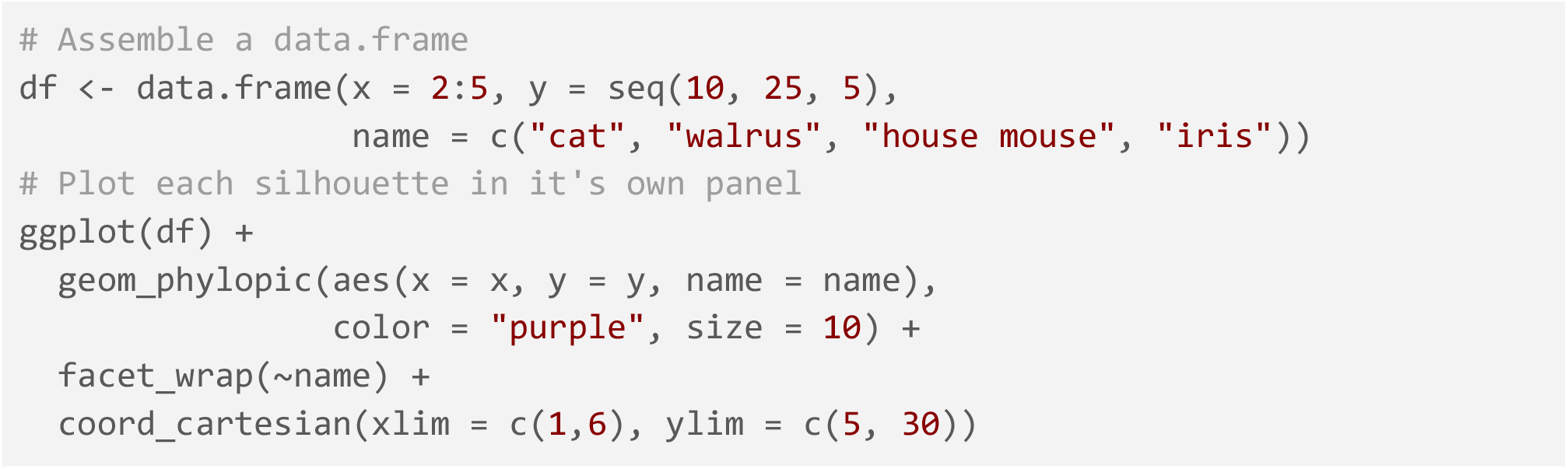

As mentioned above, for convenience these three functions all allow for transformations during the plotting stage. For the `add_phylopic` and `add_phylopic_base` functions, `horizontal`, `vertical`, `alpha`, and `color` are arguments. For the `geom_phylopic` function, these are ggplot2 aesthetics.

### Attribution

The rphylopic package gathers resources (i.e. silhouettes) from the PhyloPic database. The silhouettes are made available under various Creative Common licenses and should be appropriately attributed when used (see figure captions for examples). To help facilitate this process, we provide the function `get_attribution`:

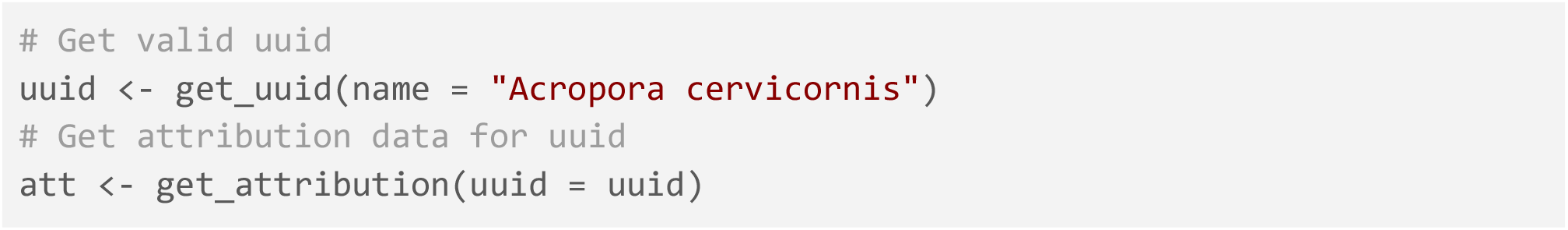

This function returns attribution data for a specific image UUID, including: contributor name, contributor UUID, contributor contact, image UUID, and the type of copyright license. While not requested by the creator of PhyloPic (T. Michael Keesey), we also encourage acknowledgement of his work.

If you use the rphylopic package in your workflow, we ask that you also cite the package appropriately. A citation for the package can be accessed via the function `citation`:

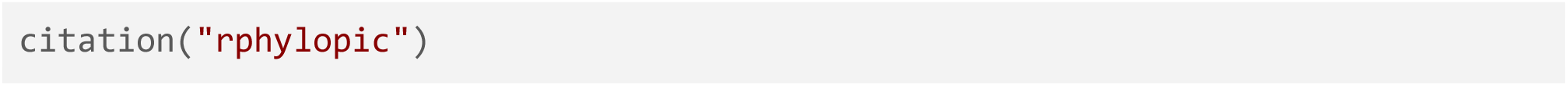

## Application

Herein we provide three example applications of the rphylopic package in combination with ggplot2 (Wickham, 2016). However, we note that all demonstrated functionality is also available for base R and showcased in the associated vignette of the package.

### Basic Accession and Transformation

The rphylopic package provides robust and flexible tools to access and transform PhyloPic silhouettes. Here we demonstrate this using the example dataset of Antarctic penguins from the palmerpenguins R package (Horst et al., 2020). Since we do not know which silhouette we want to use, we first use `pick_phylopic` to identify a suitable silhouette for *Pygoscelis*. We then rotate this silhouette and use it instead of points for a standard x-y plot showing the relationship between bill length and flipper length (Fig. 2). Note that we modify the size and colour of the silhouette based on the mass and sex of each penguin, respectively. As with all ggplot-related functions, `geom_phylopic` accepts both ‘color’ and ‘colour’ as aesthetic names.

**Figure 2.**
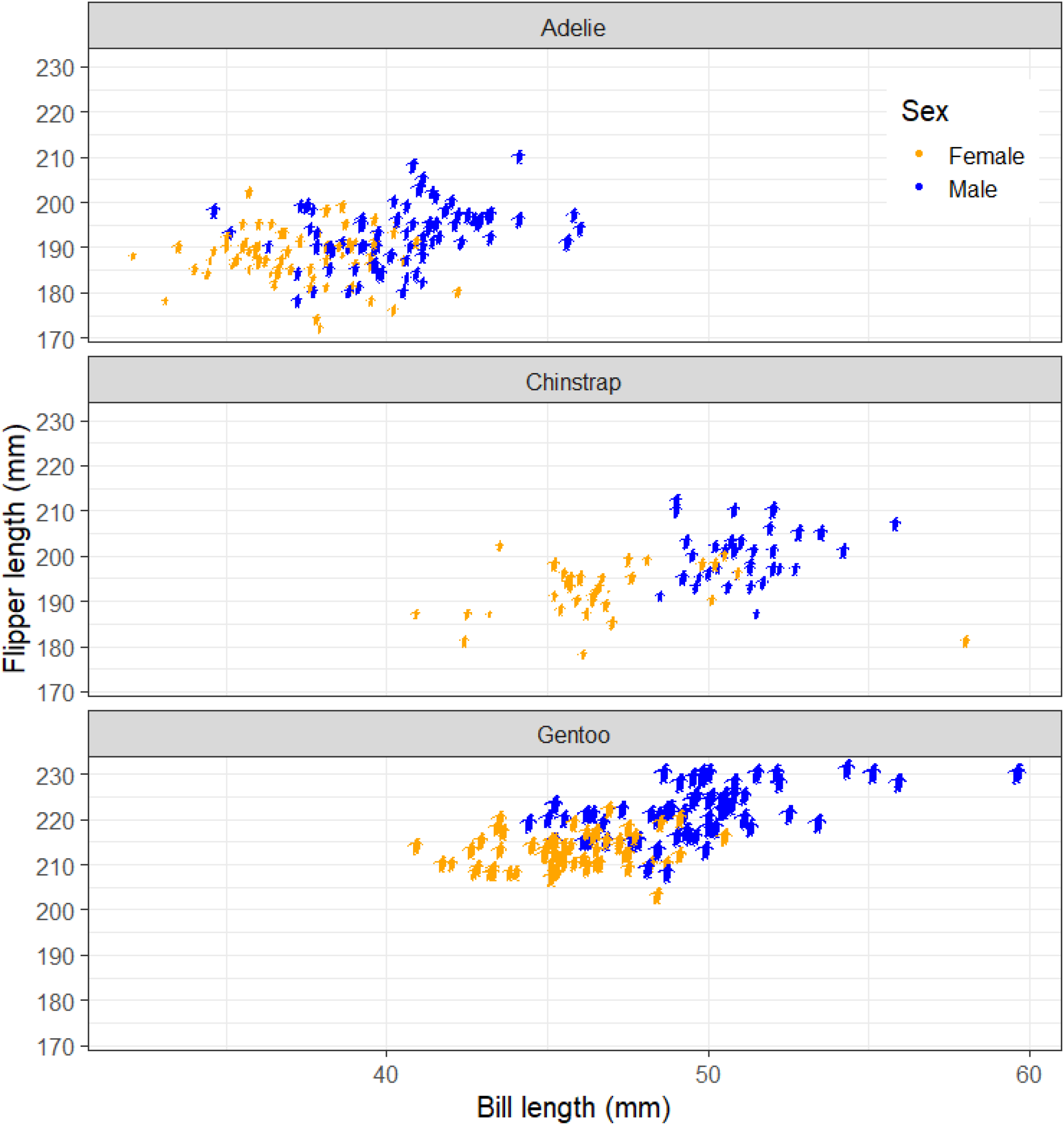
The relationship between bill length and flipper length in Antarctic penguins. Each panel represents a different species of *Pygoscelis*. Silhouette sizes represent relative body masses and silhouette colours represent the sex of measured individuals (orange: female; blue: male). The image silhouette is from PhyloPic (https://www.phylopic.org/; T. Michael Keesey, 2023) and was contributed by Alexandre Vong (2018; CC0 1.0).

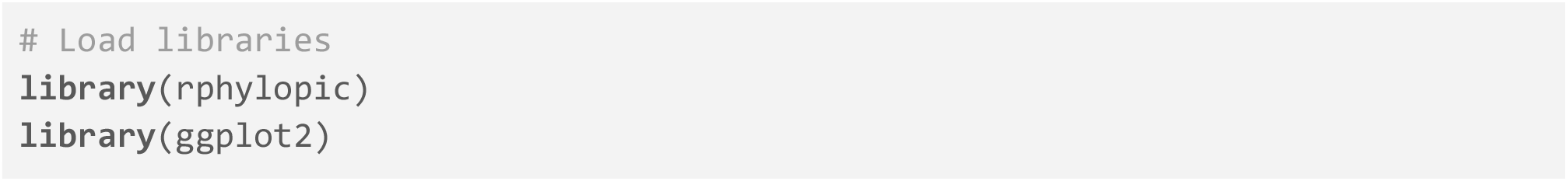
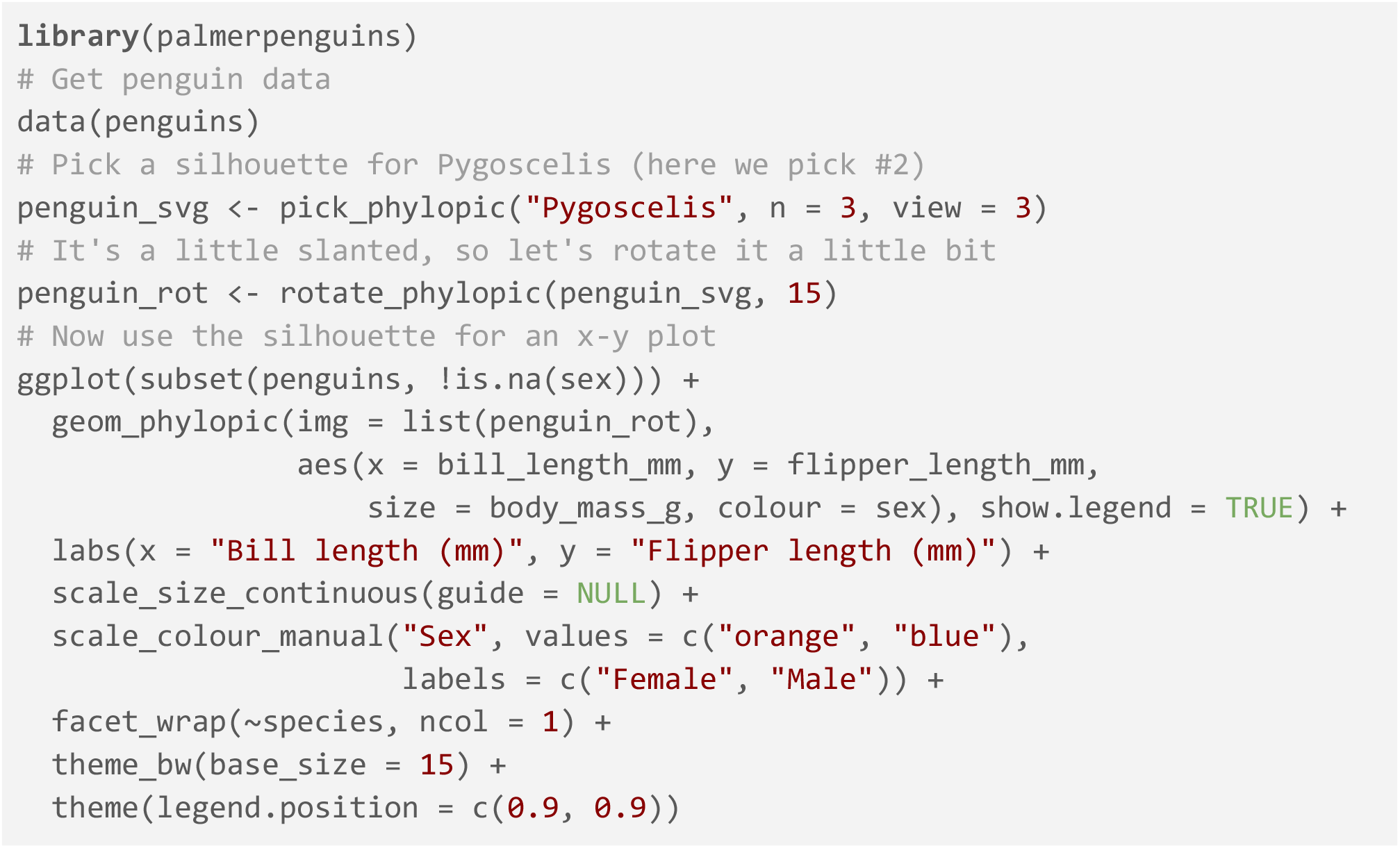

### Geographic distribution

In much the same way as generic x-y plotting, the rphylopic package can be used in combination with ggplot2 (Wickham, 2016) to plot organism silhouettes on a map (Fig. 3). That is, to plot data points (e.g. species occurrences) as silhouettes. We provide an example here of how this might be achieved for individuals who wish to do so. For this application, we use the example occurrence dataset of early (Carboniferous to Early Triassic) tetrapods from the palaeoverse R package (Jones et al., 2023) to visualise the geographic distribution of *Mesosaurus* fossils.

**Figure 3.**
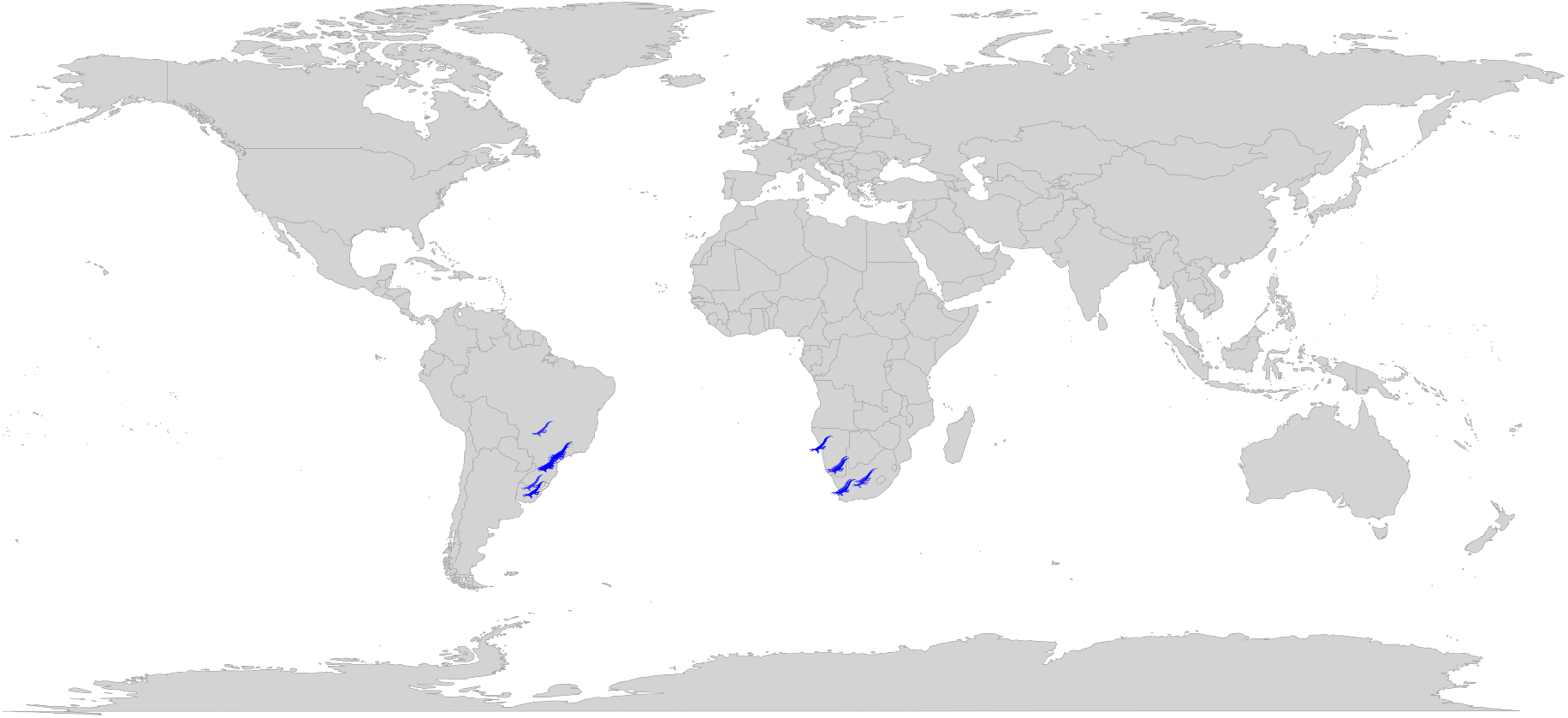
The geographic distribution of Carboniferous to Early Triassic *Mesosaurus* fossil occurrences. The image silhouette is from PhyloPic (https://www.phylopic.org/; T. Michael Keesey, 2023) and was contributed by Antoine Verrière (2016; CC BY-NC-SA 3.0). The original silhouette has been modified in colour from black to blue.

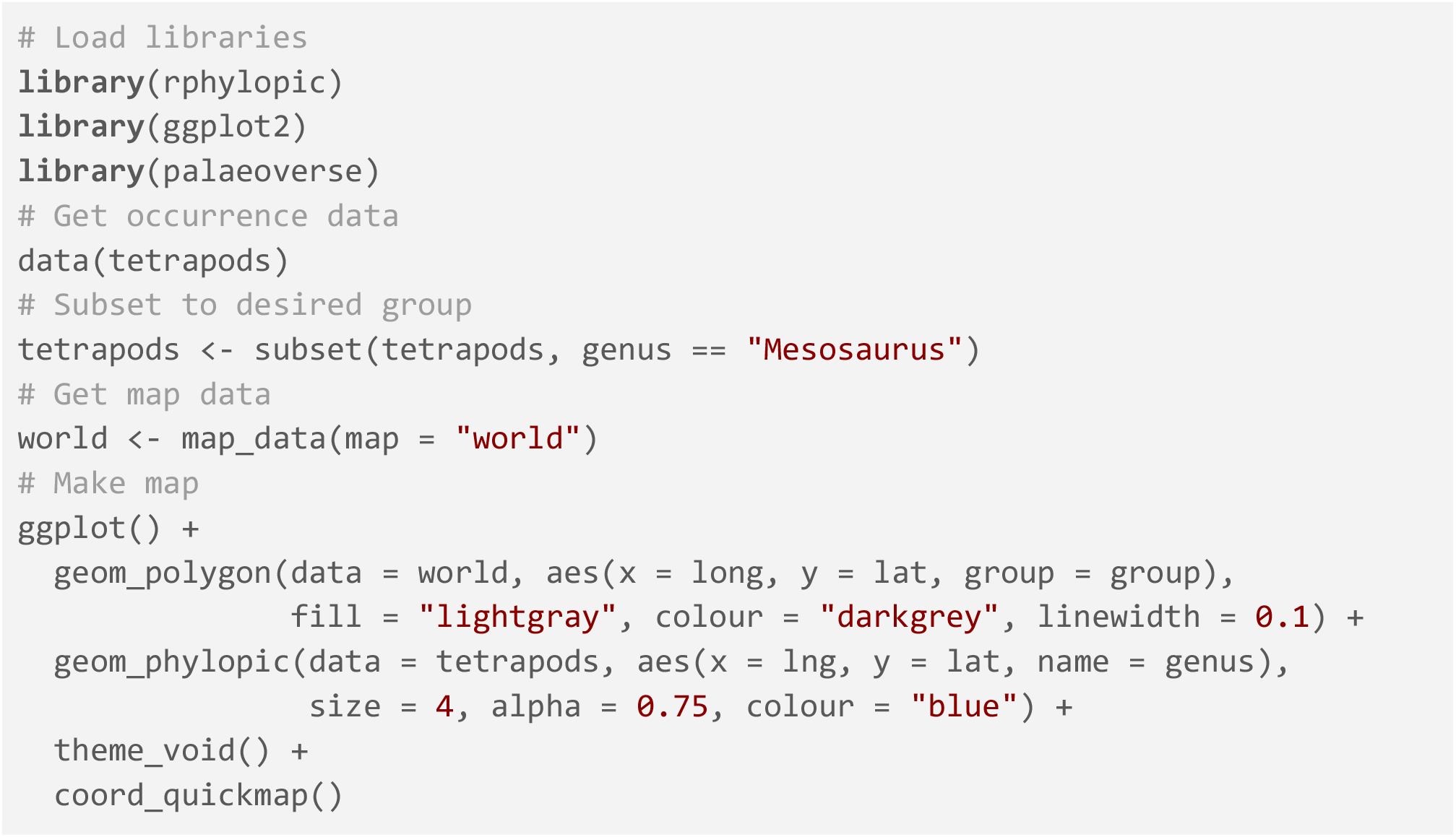

### Phylogenetics

Another common use case of PhyloPic silhouettes is to represent taxonomic information. In this example, we demonstrate how to use silhouettes within a phylogenetic framework (Fig. 4). In this case, the phylogeny includes taxa across all vertebrates. Even many taxonomic experts are unlikely to know the scientific names of these 11 disparate taxa, so we replace the names with PhyloPic silhouettes. We use the ggtree R package (Yu et al., 2017) to plot the phylogeny and the deeptime R package (Gearty, 2023) to add a geological timescale to the background. We use a vectorized version of the `get_uuid` function to retrieve UUID values for all of the species at once; however, one of the scientific names is not matched in the PhyloPic database so we need to use a higher taxonomic name. We also choose to use the `pick_phylopic` function to select a specific image for the boar. Note that only a single size is specified and aspect ratio is always maintained, hence why the silhouettes all have the same height but different widths.

**Figure 4.**
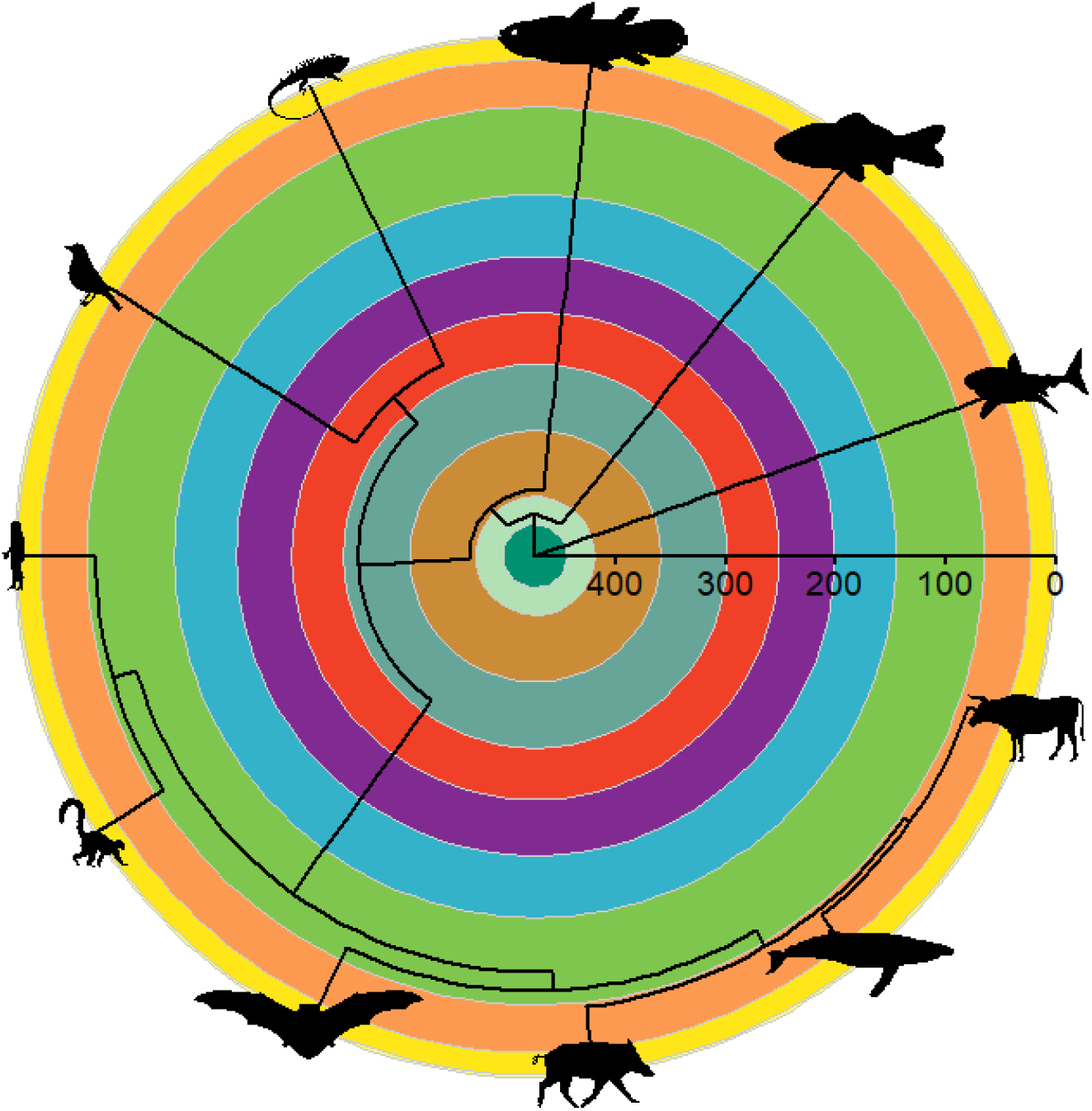
A very simplified phylogeny showing the relationships between a selection of extant vertebrates. The coloured background represents geological periods as defined by the International Commission on Stratigraphy, and the axis represents time in millions of years before present. The image silhouettes are from PhyloPic (https://www.phylopic.org/; T. Michael Keesey, 2023) and were contributed as follows: *Carcharodon carcharias* by Margot Michaud (2018; CC0 1.0); *Carassius auratus* by Corrine Avidan (2022; CC0 1.0); *Latimeria chalumnae* by Chuanxin Yu (2021; CC0 1.0); *Homo sapiens* by T. Michael Keesey (2011; CC0 1.0); *Lemur catta* by Ferran Sayoi (2015; CC0 1.0); *Lasiurus cinereus* by Andy Wilson (2022; CC0 1.0); *Sus scrofa* by Ferran Sayoi (2015; CC0 1.0); *Megaptera novaeangliae* by T. Michael Keesey (2011; CC BY-SA 3.0); *Bos taurus* by T. Michael Keesey (2011; CC BY-SA 3.0); *Iguana iguana* by Jack Mayer Wood (2014; CC0 1.0); and *Turdus migratorius* by Karina Garcia (2022; CC BY-NC 3.0).

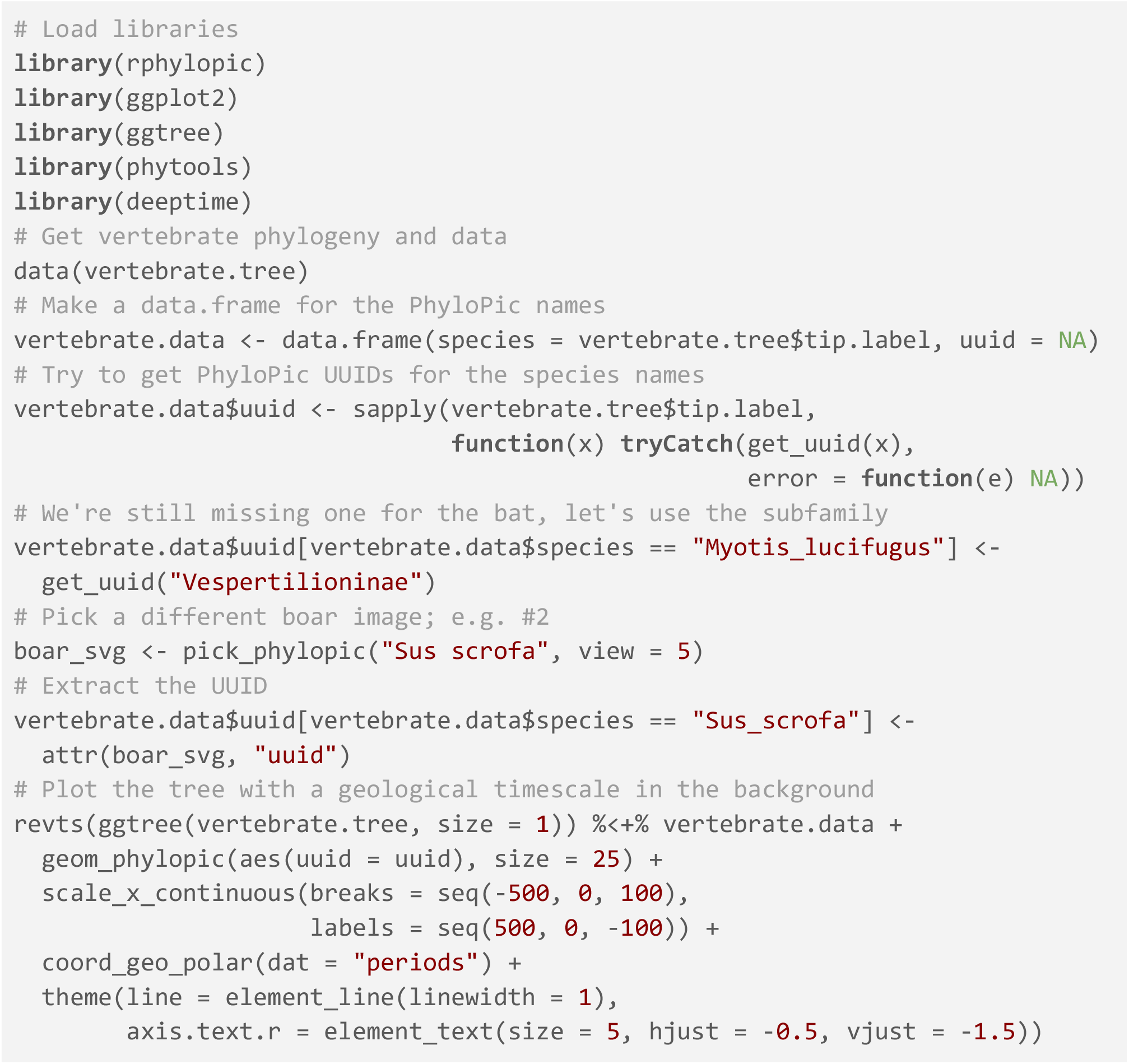

## Resources

We have made several resources available for our users. First, we have built a package website (http://rphylopic.palaeoverse.org) which provides information on how to contribute to rphylopic, report issues and bugs, and a contributor code of conduct. We have also made available a cheat sheet and vignette (tutorial) for the package which provides a user-friendly usage guide (see package website).

## Availability

The rphylopic R package can be installed via the Comprehensive R Archive Network (CRAN) using the `install.packages(“rphylopic”)` function. The package has been tested on Windows, Mac, and Linux (Ubuntu) operating systems using GitHub actions. Source code for the package is available at https://github.com/palaeoverse-community/rphylopic.

## Acknowledgements

We thank Scott Chamberlain, the developer and maintainer of earlier versions of rphylopic (prior to ver. 1.0.0). While the package has been overhauled by the current authors, we would be amiss to not acknowledge Scott’s initial efforts. We would also like to extend our gratitude to T. Michael Keesey for the PhyloPic project and all those who contribute silhouettes to PhyloPic. Without such efforts, rphylopic would simply not exist. Finally, we would like to thank Miranta Kouvari for producing the logo and cheat sheet for the rphylopic package. The contributions of W.G. were supported by the Lerner-Gray Postdoctoral Research Fellowship from the Richard Gilder Graduate School at the American Museum of Natural History. The contributions of L.A.J. were supported by a Juan de la Cierva-formación 2021 fellowship (FJC2021-046695-I / MCIN / AEI / 10.13039 / 501100011033) from the European Union “NextGenerationEU”/PRTR. L.A.J. also received funding from the European Research Council under the European Union’s Horizon 2020 research and innovation program (grant agreement 947921; MAPAS project).

## Notes

https://github.com/palaeoverse-community/rphylopic

### Competing Interest Statement

The authors have declared no competing interest.

